# A statistical modelling reveals bi-directional chromatin scanning by RAG in the human TCR-α locus

**DOI:** 10.1101/2020.02.21.959197

**Authors:** Ciarán O’Connor, Deirdre McNamara, Shigeki Nakagome

**Affiliations:** School of Genetics & Microbiology, Trinity College Dublin, Dublin, Ireland; Trinity Academic Gastroenterology Group, Department of Clinical Medicine Tallaght Hospital, Trinity College Dublin, Dublin, Ireland; School of Medicine, Trinity College Dublin, Dublin, Ireland; Trinity Translational Medicine Institute, Trinity College Dublin, Dublin, Ireland

**Author notes:** **Correspondence**: Shigeki Nakagome, **Address:** Room 0.79, Trinity Translational Medicine Institute, St. James’s Hospital, James’s Street, Dublin 8, Ireland, **E-mail:**.

## Abstract

Rearrangements of germline variable (V), diversity (D), and joining (J) coding gene segments generate a diverse T cell receptor (TCR) repertoire. Two different molecular mechanisms govern a stochastic nature of gene choices by the recombination activating gene (RAG) protein complex. One is diffusion access in which a chromatin loop containing a coding gene bounces back and forth until the loop encounters a distant gene. The other is a linear chromatin scan, which extrudes chromatin loops to assemble a gene pair in a distance-dependent manner. However, the extent to which those two mechanisms underlie V(D)J recombination in human TCRs still remains unclear. Applying statistical modelling to TCR sequence data, we infer the stochastic process of gene choices during this recombination. The inferred usage of V and J gene segments vary from gene to gene. This pattern of gene usage is consistent across individuals, suggesting a fundamental bias in the gene choice. Our modelling shows the dependency of VJ pairing on their linear distance in the TCR-α locus; proximal-proximal and distal-distal VJ pairs are more preferred than proximal-distal or distal-proximal gene pairs. These results support bi-directional RAG scanning into proximal and distal sides of the TCR-α V gene locus.

## Introduction

T cell receptors (TCRs) expressed on the T cell surface mediate the recognition of peptide antigen presented by major histocompatibility complex (Hedrick et al., 1984; Yanagi et al., 1984). The germline structure of the TCR loci (α-chain on chromosome 14, TCR-α; β-chain on chromosome 7, TCR-β) is characterized by paralogous gene segments for the variable (V), diversity (D), and joining (J) coding genes (see Figure 1). During T cell development in the thymus, somatic recombination proceeds by selecting one of each type of coding gene segments and joining V and J gene segments for TCR-α or V, D, and J gene segments for TCR-β, with random nucleotide additions or deletions at the junctions. This random shuffling, so-called V(D)J recombination, is vital for producing highly diverse sequence repertoires.

**Figure 1.**
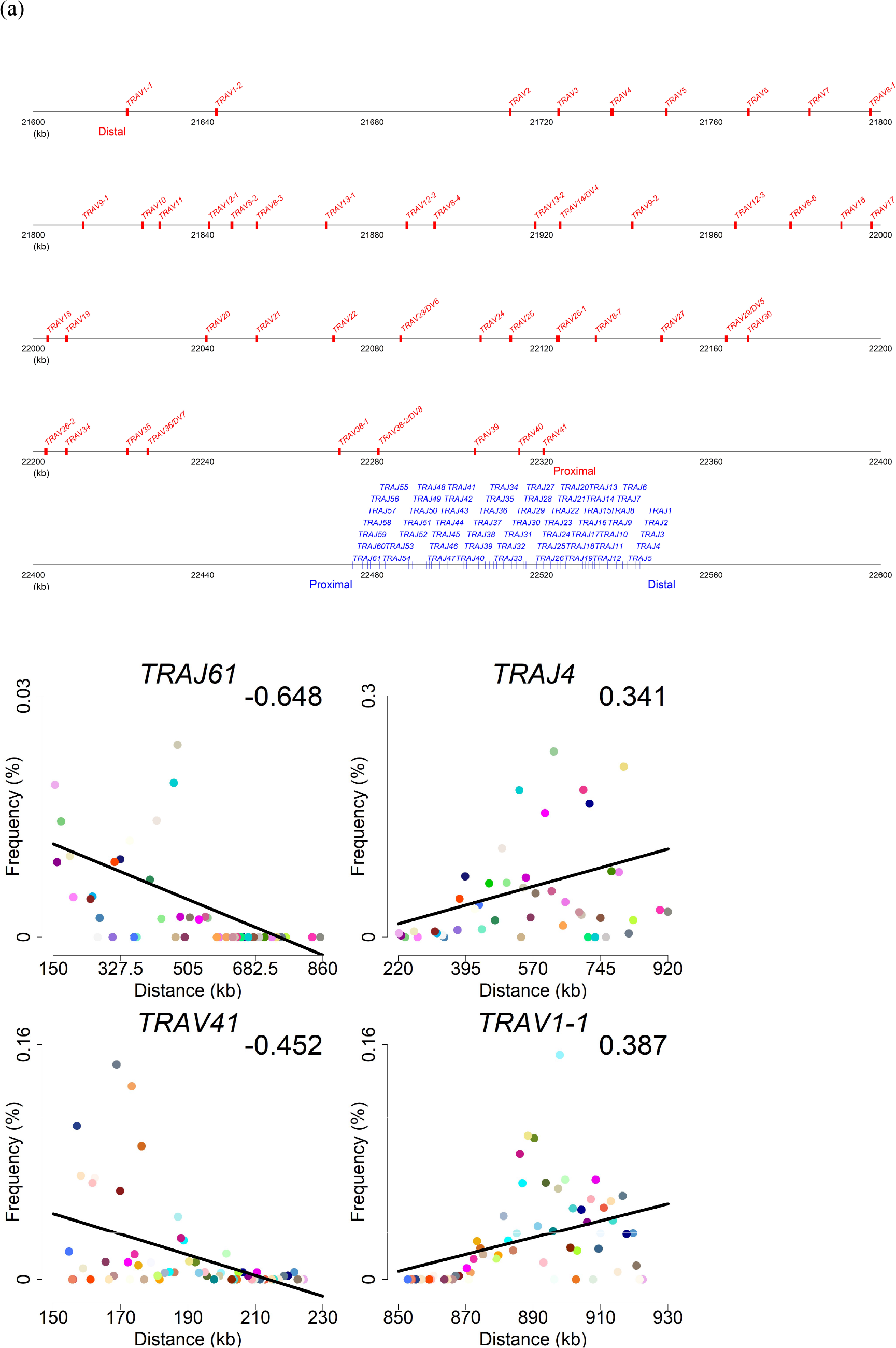
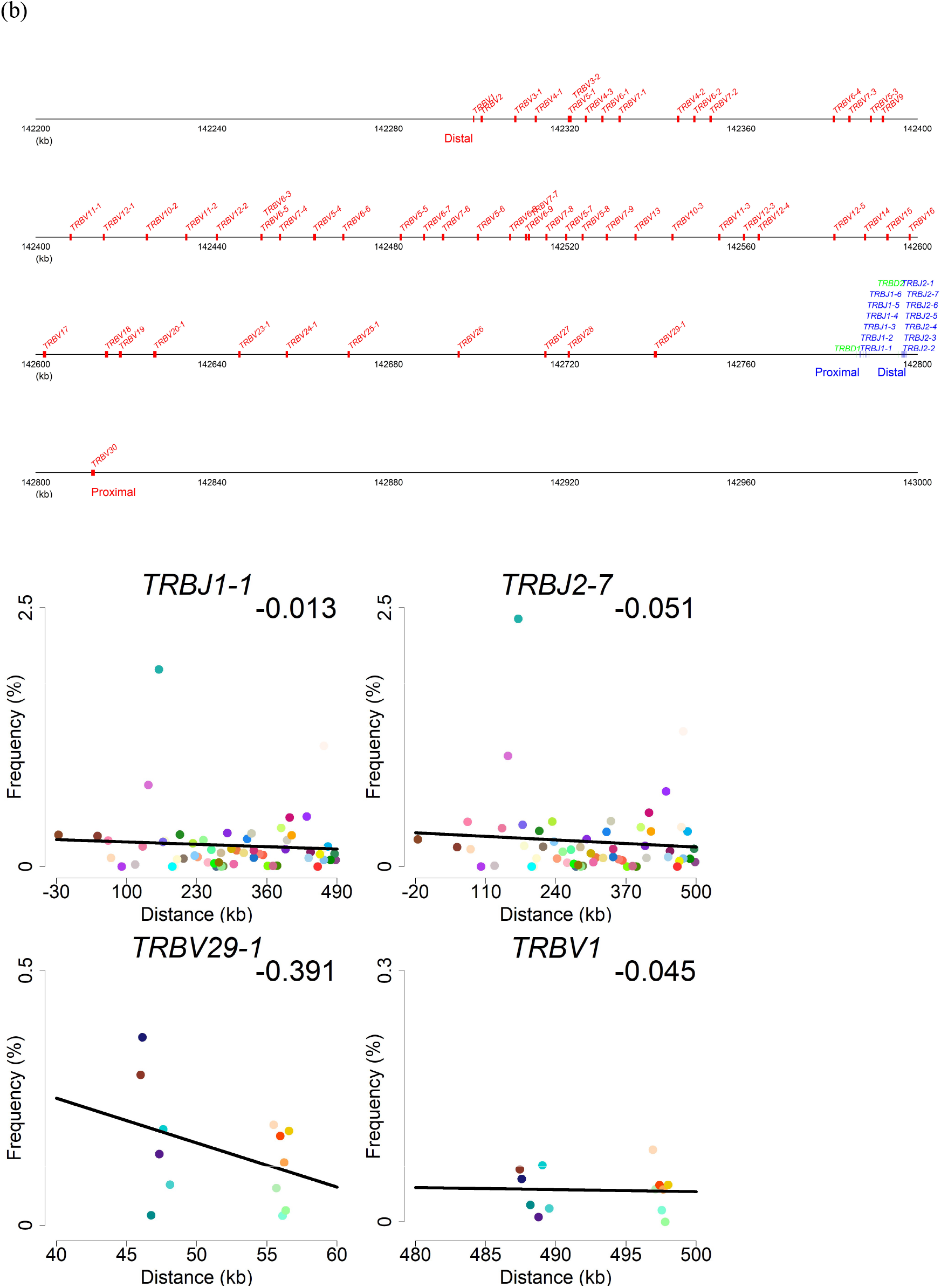
Germline structures and frequencies of gene pairs between a given V gene segment and all J gene segments or between a given J gene segment and all V gene segments in the (a) TCR-α and (b) TCR-β loci. Four gene segments are selected for each locus as examples. A full set of gene segments is shown in Supplementary Figs. 4-5 (TCR-α) and Supplementary Figs. 6-7 (TCR-β). Different colours in each plot represent different V or J gene segments. The positive or negative value in the plot shows a correlation coefficient (*rho*) between the gene usage and the distance from a given gene segment.

V(D)J recombination is initiated by the recombination activating gene (RAG) 1 and 2 proteins, which form a heterotetramer (Kim et al., 2015) and recognize recombination signal sequences (RSSs) flanking each of the coding gene segments (Schatz and Swanson, 2011). Upon binding to RSSs in a given pair of the gene segments, the RAG complex introduces DNA double-stranded breaks between the gene segments and their RSSs (Swanson, 2004; Puebla-Osorio and Zhu, 2008; Helmink and Sleckman, 2012). Those cleaved gene ends then undergo nucleotide deletions and non-templated nucleotide additions before being re-joined to form a TCR. The combinatorial variation of different coding gene segments, coupled with the junctional variation, gives rise to 10^15^~10^21^ potential TCRs (Davis and Bjorkman, 1988).

Growing evidence from research on the mouse immunoglobulin heavy chain (*Igh*) locus, where V, D, and J gene segments are rearranged during B cell development, demonstrates that the RAG complex employs two different molecular mechanisms to search for a pair of the coding gene segments: diffusion access (Lucas et al., 2014) and linear chromatin scanning (Hu et al., 2015; Jain et al., 2018). Locus contraction brings V gene segments that are linearly distant into physical proximity of a rearranged DJ segment (Kosak et al., 2002; Fuxa et al., 2004; Degner et al., 2011; Guo et al., 2011; Medvedovic et al., 2013; Bolland et al., 2016; Montefiori et al., 2016; Johanson et al., 2019). In the diffusion access, the V gene segment most likely to be chosen for a DJ segment depends on how many times each V segment encounters the DJ segment while a loop containing the DJ segment is bounced back and forth in a spring-like fashion (Lucas et al., 2014). On the other hand, the RAG complex can scan chromatin by extruding the loops and drawing distal chromosomal regions into physical proximity in a linear fashion (Vietri Rudan et al., 2015; Vian et al., 2018). However, the extent to which those two different mechanisms underlie random pairing in the human TCR loci is largely unknown mainly due to a technical difficulty in generating data from developing T cells in the human thymus. Here, we apply a quantitative statistical method (Murugan et al., 2012; Marcou et al., 2018) to model the stochastic process of gene choices during V(D)J recombination using bulk sequence data of TCR-α and -β repertoires.

## Materials and Methods

### Quality control for TCR sequencing data

We used bulk sequence data of TCR-α and -β chain repertoires that were generated from peripheral blood of 11 individuals in a previous study (Saravanarajan et al., 2020). Raw sequence reads were pre-processed with the MiXCR version 3.0.2 (Bolotin et al., 2015, 2017) to merge paired-end reads, correct for PCR and sequencing errors, and remove reads that are low quality or do not fully cover complementarity-determining region 3 (CDR3) region. We only used sequences that were non-productive due to out-of-frames or stop codons in the CDR3 region (*i.e.* on average 10.68% of total reads) for subsequent statistical modelling.

### Probabilistic modelling on V(D)J recombination

We applied the IGoR software (Marcou et al., 2018) to the unique sequences of non-productive rearrangements to model the recombination process without impacts of thymic selection. The sequences were aligned against 103 V genes (*TRAV*) and 68 J genes (*TRAJ*) for TCR-α chains or 89 V genes (*TRBV*), 3 D genes (*TRBD*), and 15 J genes (*TRBJ*) for TCR-β chains with default settings of parameters. We then estimated parameters of the recombination models where the observed sequences result from different stochastic events including the choice of germline genes, deletions, and insertions. Once the models were inferred, we simulated 100,000 sequences for the TCR-α and -β chains respectively and calculated the usage of all V and J gene pairs (7,004 pairs for TCR-α and 1,335 pairs for TCR-β). By repeating this simulation step 100 times, we measured variation in the gene usage across simulated data (Supplementary Fig. 1).

### Correlation analysis of paired gene usage between individuals

To assess variation in the inferred models across individuals, we measured Spearman’s rank correlation coefficients (*rho*) in the usage of gene pairs between all pairwise individuals. There are 7,004 pairs (103 *TRAV* and 68 *TRAJ*) and 1,335 pairs (89 *TRBV* and 15 *TRBJ*) at the TCR-α and TCR-β loci, each of which has the frequency in a given individual (Supplementary Fig. 2).

### Dependency of paired gene usage on the physical distance

We tested whether the choice of V and J gene pairs is dependent on the physical distance between those two genes. Start and end positions of each gene were retrieved from the human reference genome (GRCh38.p13) using the Ensembl Genome Browser (version 97) (Zerbino et al., 2018) (Figure 1). We then tested the significance of correlations between the distances and the usage of gene pairs (*i.e.* a given V gene and all J genes or a given J gene and all V genes) in a positive or negative direction by using the Spearman’s *rho* (Figure 1; Supplementary Figs. 3-6). Since some genes have multiple alleles, we took a sum of the paired gene usage over those alleles. Multiple testing was corrected with the Benjamini & Hochberg method (Benjamini and Hochberg, 1995).

To identify patterns in the shift of preference as to the gene choice by V or J genes, we tested the linearity of relationships between the *rho* values and the median distances from each V to all J genes or from each J gene to all V genes (Supplementary Figs. 7-8). The significance of linear relationships was calculated for all individuals (Figure 2; Supplementary Fig. 9).

**Figure 2.**
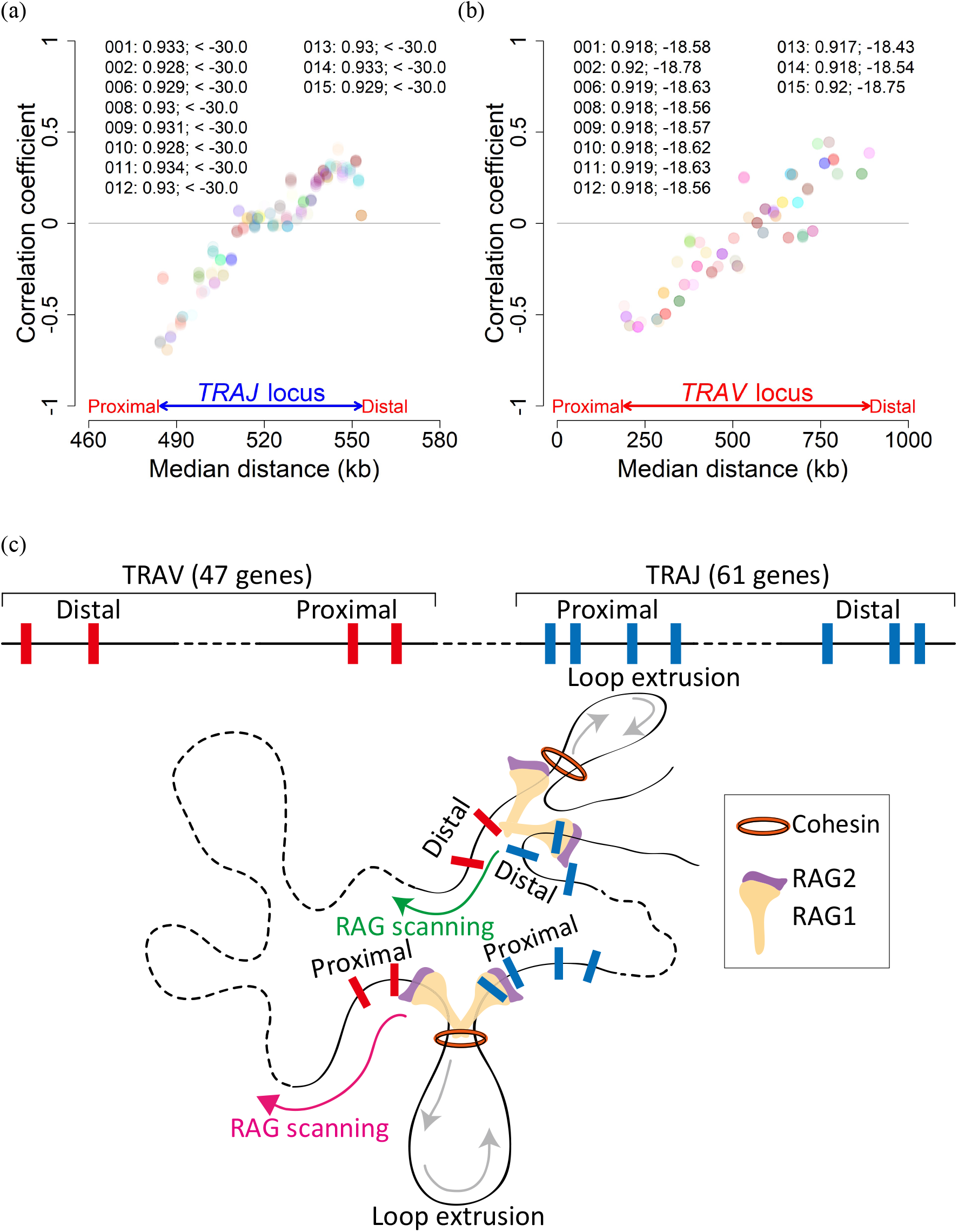
Plots of correlation coefficients (*rho*; Supplementary Figs. 4-5 and 8) against median distances (a) between each *TRAJ* and all *TRAV* segments or (b) between each *TRAV* and all *TRAJ* segments. Different colours show (a) different *TRAJ* or (b) different *TRAV* genes, overlaid across al individuals. The linearity of these plots is tested for each individual; the values show correlation coefficients (*rho*) and log_10_-scaled *p*-values. (c) A recombination centre, where RAG proteins and transcriptional factors are recruited, is created at the most proximal *TRAJ* segment. The RAG complex can then search for a *TRAV* partner through cohesin-mediated loop extrusion. Given that the RAG complex prefers to choose genes first encountered, this scanning is likely to result in the decrease of the *TRAV* usage over the distance from the recombination centre. On the other hand, another RAG complex is recruited into the most distal *TRAJ* segment, which scans and utilizes distal *TRAV* genes through loop extrusion.

### Analysis of open chromatin regions

We used Assay for Transposase-Accessible Chromatin using sequencing (ATAC-seq) data generated from fetal thymocytes (Calderon et al., 2019) to measure changes of chromatin accessibility in the TCR-α locus during T cell development. We applied a stringent cut-off of FDR<1% to define accessible genomic regions in four different thymocytes (*i.e.* double negative, pre-T double negative, immature single positive, and double positive cells) across three individuals (Supplementary Fig. 10).

## Results and discussion

In both the TCR-α and -β loci (Supplementary Fig. 1), the frequencies of V or J gene usage are different from genes to genes. Given that each gene has a little variation in its usage across 100 replicates, this non-uniformity is unlikely to be due to stochastic effects. Comparing the usage of all paired gene segments between individuals, we confirmed that this pattern of gene usage is consistent across individuals with *rho*>0.99 (Supplementary Fig. 2). These results suggest that V(D)J recombination may introduce mechanistic constraints on the V and J gene choices.

Synergy of diffusion access and chromatin scanning is crucial to providing the RAG complex with equal accessibility to all gene segments for pairing (Jain et al., 2018). However, if one mechanism has a stronger impact on pairing than the other, the choice of gene pairs may be biased towards specific combinations. A key factor that differentiates those two mechanisms is the accessibility to proximal V genes. Diffusion access requires locus contraction first to bring different V gene segments into physical proximity of rearranged DJ segments before starting searching for interaction partners (Lucas et al., 2014). On the other hand, the RAG complex gains access to proximal V gene segment without locus contraction by means of linear chromatin scanning (Fuxa et al., 2004; Jain et al., 2018). Therefore, if RAG chromatin scanning takes an active role in gene pairing, proximal V gene segments are likely to be utilized more frequently than distal V segments, which can be detectable as a distance-dependent utilization pattern.

To test this hypothesis, we focused on how frequently each J gene segment makes pairs with different V gene segments and correlated those frequencies with their distances. In the TCR-α locus, the *TRAV* usage decreases over the distance from a given *TRAJ* (*e.g. TRAJ61*), as expected from the distance-dependency of gene choices (false discovery rate, FDR, <10%; Figure 1a; Supplementary Fig. 3). However, we only observed this negative correlation when *TRAJ* genes were linearly close to the *TRAV* locus (*i.e.* 5’-end of the *TRAJ* locus). Even though the *TRAJ* locus spans a relatively short distance, <70 kb, the direction of correlation relationships turns out to be positive in *TRAJ* genes (FDR<10%) that are distal to the *TRAV* locus (*i.e.* 3’-end of the *TRAJ* locus; *e.g. TRAJ4* in Figure 1a). These results suggest that the choice of gene segments for pairing may depend on where *TRAV* and *TRAJ* segments are linearly located on the chromosome.

We then tested whether the same pattern is still observable in the comparison between the usage of each V gene across all J genes and their distances (Figure 1a; Supplementary Fig. 4). Proximal *TRAV* segments (*e.g. TRAV41*) tend to be chosen by proximal *TRAJ* more often than distal *TRAJ*, supporting negative correlations (FDR<10%). In contrast, distal *TRAV* segments (*e.g. TRAV1-1*) are preferred by distal *TRAJ* rather than proximal *TRAJ* (FDR<10%). Therefore, the preference for the gene choice is shaped by bi-directional dependency of proximal *TRAV* and proximal *TRAJ* genes or distal *TRAV* and distal *TRAJ* genes. None of the correlations in the TCR-β locus supported distance-dependency (Figure 1b; Supplementary Figs. 5-6), implying that synergy of the gene choice mechanisms may be more balanced in the TCR-β locus than in the TCR-α locus.

Plotting a correlation coefficient against a median distance between a given *TRAJ* and all *TRAV* or between a given *TRAV* and all *TRAJ* (Supplementary Fig. 7), we found that the more proximal or distal *TRAJ* and *TRAV* segments are, the more significant correlations become (FDR<10%). Furthermore, the correlation is continuously shifting from the negative to the positive direction along the *TRAJ* or *TRAV* locus. This shift is significantly linear with *rho*>0.91, which is consistent across all individuals (Figures 2a-b). We also confirmed that this linear relationship is absent in the TCR-β locus and unique to the TCR-α locus (Supplementary Figs. 8-9). Upon the recruitment of RAG proteins and transcriptional factors to a recombination centre at the *TRAJ* locus (Ji et al., 2010), the RAG complex can initiate its scanning towards the *TRAV* locus (Jain et al., 2018). Indeed, the *TRAJ* locus becomes more accessible at the stage of double positive cells, where rearrangements of the TCR-α chains occur, than that of double negative cells or immature single positive cells in the foetal thymus (Supplementary Fig. 10). This pattern of chromatin accessibility supports the recruitment of the recombination centre into both proximal and distal *TRAJ* segments. However, the shift in the direction of correlation relationships cannot be explained by RAG scanning into proximal *TRAV* as such scanning is unlikely to get access to distal *TRAV* and increase their usage while ignoring all proximal *TRAV* passed through. Rather, the RAG complex may be able to begin to scan the most proximal and most distal *TRAV* segments respectively, as proposed in the mouse *Igh* locus (Jain et al., 2018), and to utilize the genes for recombination by giving preference to the ones first encountered (Figure 2c).

In conclusion, this study applies the quantitative statistical method to modelling of V(D)J recombination in human TCRs and provides evidence that RAG chromatin scanning plays a critical role in pairing *TRAV* and *TRAJ* segments. With increasing scale and complexity of TCR sequence data, statistical modelling becomes a powerful tool for advancing our understanding of TCR biology including the recombination process (DeWitt et al., 2018; Venturi and Thomas, 2018; Bradley and Thomas, 2019; Dupic et al., 2019). Given that the models of TCR recombination assumed in this study are still simplified mechanistically (Murugan et al., 2012; Marcou et al., 2018), however, our findings need to be further validated with experimental approaches. Such an interdisciplinary effort will make it possible to test hypotheses and improve models in more efficient and interactive ways than ever as seen in emerging fields of immunogenomics (Zewde et al., 2018) and systems immunology (Venturi and Thomas, 2018).

## Acknowledgements

We thank Drs. Derek Doherty, Aideen Long, Ross McManus, and Aiden Corvin for helpful comments and suggestions on the manuscript. This study was supported by grants from institutional support including Wellcome Trust Institutional Strategic Support Fund to Trinity College Dublin, Trinity Translational Medicine Institute Collaborative Pilot Study Awards, and Dean’s Research Initiatives Fund 2016-17.

